## Supplemental Figures for "Performance of localization prediction algorithms decreases rapidly with the evolutionary distance to the training set increasing"

S1 Experimentally verified and predicted organelle proteins as a percentage of the whole genome

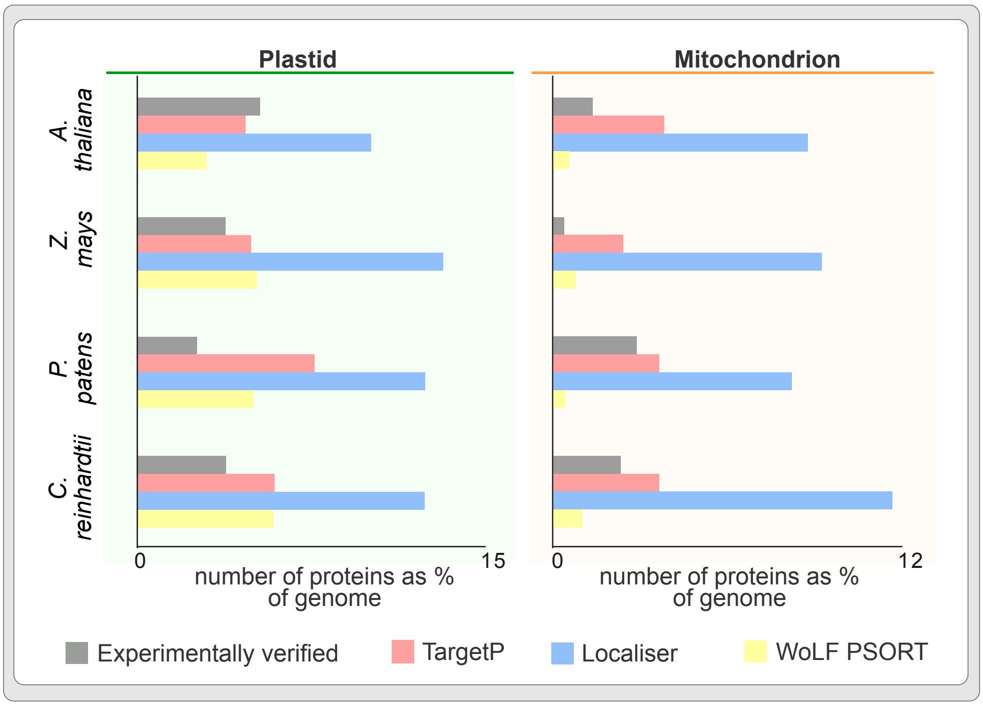


Fig. S1 Experimentally verified plastid (on the left) and mitochondrial (on the right) proteins as a percentage of all proteins encoded by a given species.

S2 Number of proteins predicted between plastid and mitochondria

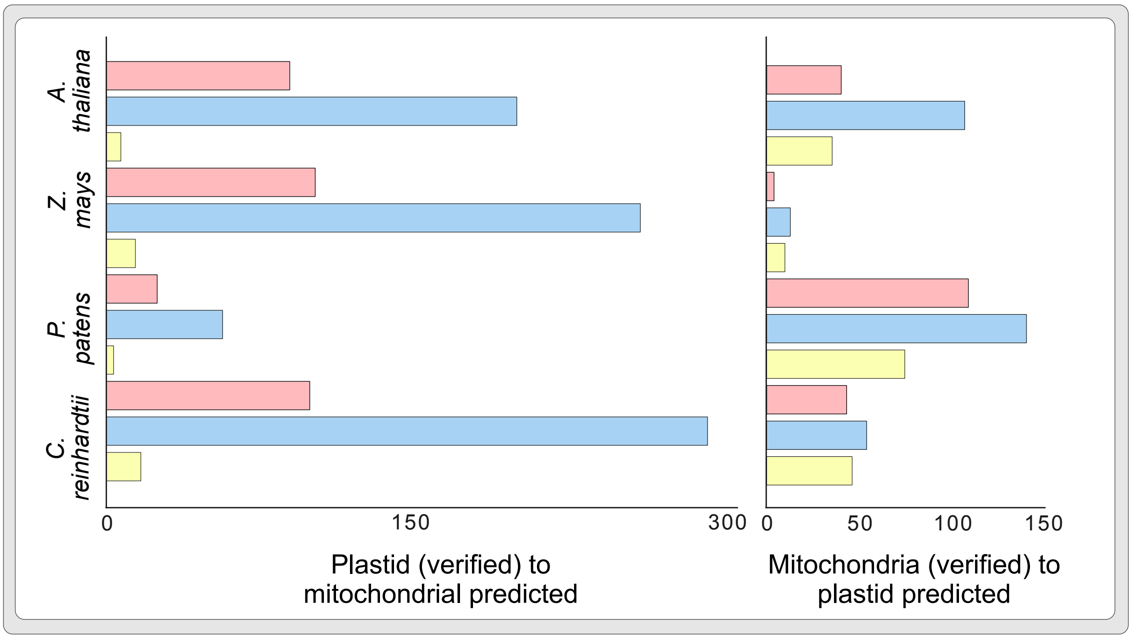


Fig. S2 Cross-organelle protein localization predictions. Experimentally verified plastid proteins that got predicted as mitochondria proteins (on the left) and verified mitochondrial proteins that got predicted as plastid proteins (on the right).

S3 Protein clustering and filtering of organelle protein families

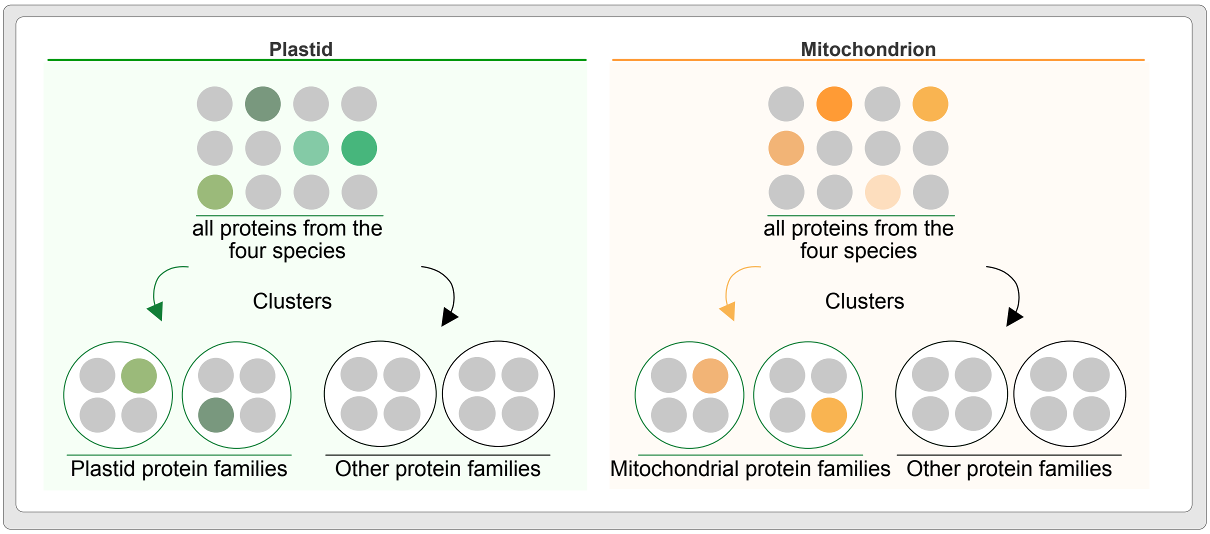


Fig. S3 Clustering of all proteins from the four photosynthetic eukaryotes and sorting of protein clusters into organelle protein families. Each circle is a protein from a species. In the first step (top boxes), source protein sequences from available species were clustered into protein families. If a protein family consisted of an experimentally verified plastid protein (in green, on the left) or a mitochondrial protein (in orange, on the right), the protein family was sorted as a plastid or mitochondrial protein family.

Fig S4 Chloroplast predicted proteins from TargetP with thylakoid predictions included

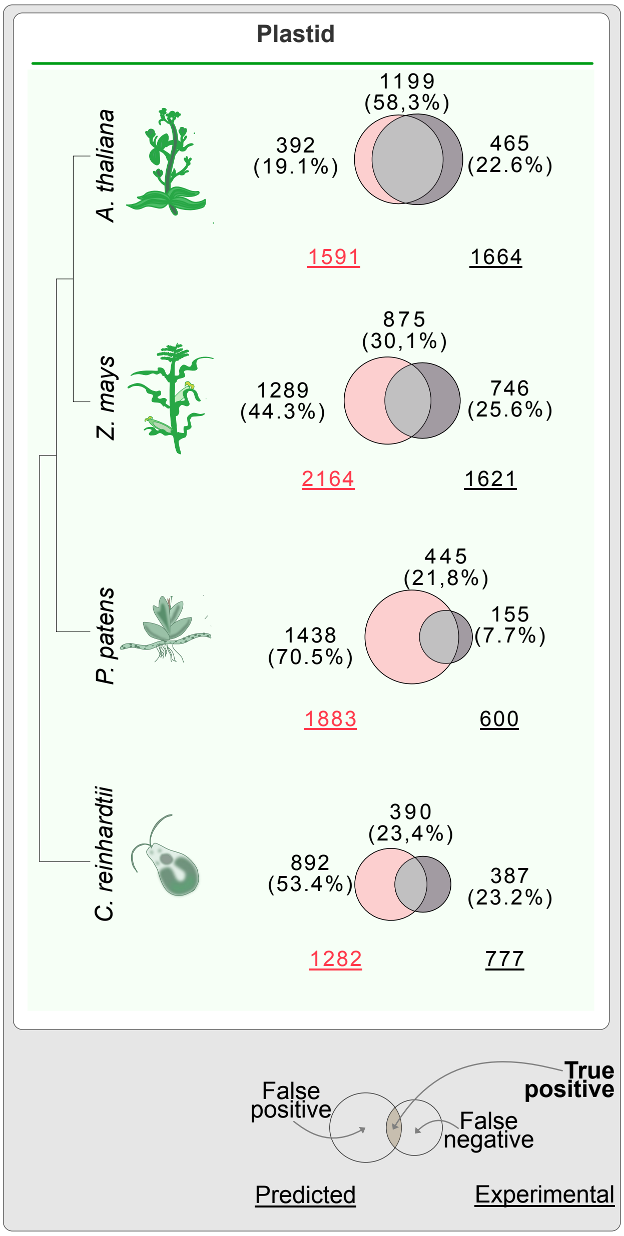


Fig. S4 Comparison of chloroplast+thylakoid proteins predicted by TargetP2.0 with experimentally localised proteins across species. Each Venn diagram represent data similar to that of Fig 1a, expect now supplemented with 'thylakoid' predicted proteins under the category 'plastid'. The Ven diagrams show an overlap between predicted (left circles) and experimentally verified organelle proteomes (right circles, grey). The underscored numbers in the bottom corners show the total number of predicted (bottom left) and experimentally confirmed proteins (bottom right). The numbers of proteins that overlap (true positives) are provided in the top right corner in bold, while the numbers of non-overlapping ones (false positives) are shown next to each circle. See also the key for the Venn diagrams on the bottom right.

Fig S5 Chloroplast predicted proteins from TargetP with thylakoid predictions included

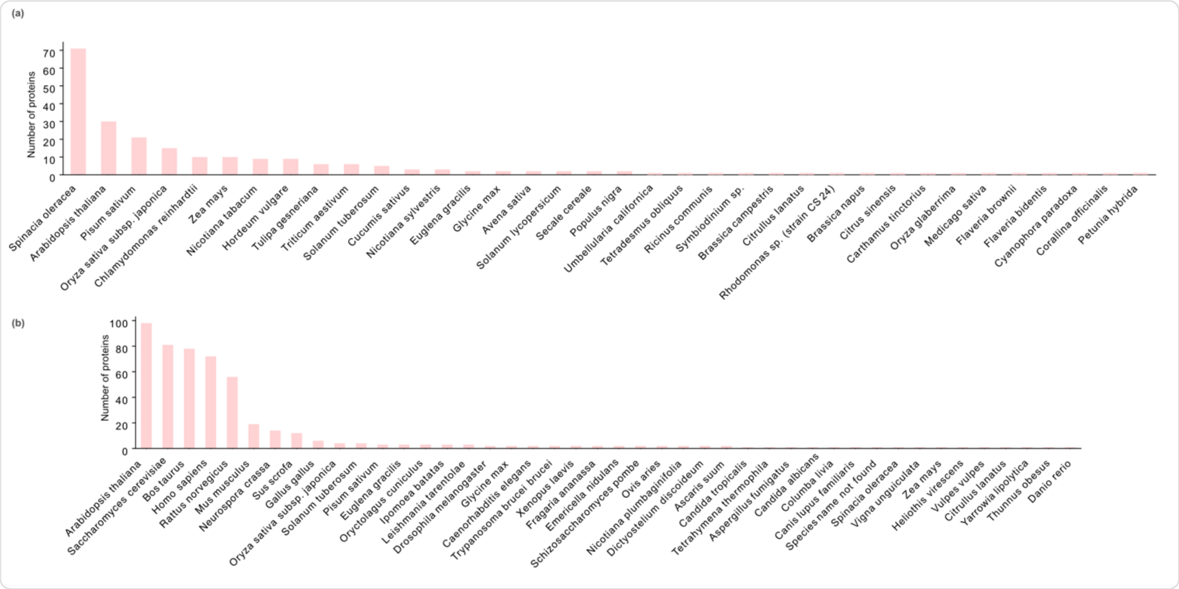


Fig. S5 Taxonomic distribution of TargetP 2.0 training dataset
